# Microglial P2X4 receptors promote ApoE degradation and cognitive deficits in Alzheimer disease

**DOI:** 10.1101/2022.05.12.491601

**Authors:** Jennifer Hua, Elvira Garcia de Paco, Nathalie Linck, Tangui Maurice, Catherine Desrumaux, Bénédicte Manoury, François Rassendren, Lauriane Ulmann

## Abstract

Numerous evidence support that microglia contributes to the progression of Alzheimer’s disease. P2X4 receptors are ATP-gated channels, which are *de novo* expressed in a subset of reactive microglia associated to various pathological contexts, contributing to microglial functions. Here, we investigated the role of P2X4 in the context of Alzheimer disease (AD). In both human AD brain and APP^swe^/PSEN1^dE9^ mice, P2X4 is almost exclusively expressed in plaque associated microglia. Genetic deletion of *P2rx4* results in the reversal of cognitive declines and in a lower amount of soluble Aß1 −42 in 12 months old APP/PS1 mice, while no obvious alteration of plaque associated microglia characteristics is observed. Using proteomic, we identified ApoE as a specific P2X4 interacting protein. We found that P2X4 regulates lysosomal cathepsin B activity promoting ApoE degradation; *P2rx4* deletion results in higher amount of intracellular and secreted ApoE in both BMDM and microglia from APP/PS1 brain. Our results support that microglial P2X4 promotes lysosomal ApoE degradation, indirectly altering Aß peptide clearance, which in turn might promote synaptic dysfunctions and cognitive deficits. Our findings also uncover a specific interplay between purinergic signaling, microglial ApoE, sAß species and cognitive deficits associated with AD.

## Introduction

Alzheimer’s disease (AD), a slowly progressive, irreversible and incurable neurodegenerative disease, is the most common form of dementia in human. The main pathological hallmarks of AD are amyloid-ß (Aß) accumulation in plaques, hyperphosphorylated Tau aggregation in neurofibrillary tangles, neuronal loss, brain atrophy and gliosis^1^. For decades, AD was mainly considered as a neuronal disease, glial cells being only considered as reacting to neuronal alterations. This neurocentric view considerably evolved in the past ten years, with both genetic and functional studies showing that neuroinflammation contributes significantly to the onset and progression of AD^2^. Indeed, genome-wide association studies (GWAS) support that approximately 50% of the susceptibility genes associated with AD are not related to neurons but to glial and vascular cells and point towards innate immune system involvement^3–5^. In the CNS, inflammation is mainly driven by two cell types, microglial cells and astrocytes. Microglia, the brain resident macrophages, are the main immunocompetent cells, which in the healthy brain, have different homeostatic functions such as monitoring neuronal activity, shaping dendritic spines, and even influencing synaptic activity^6^. In pathological conditions, microglia enter into reactive states characterized by a transcriptional and functional remodeling. Using single cell RNAseq analysis, recent studies revealed that microglial reactivity evolves along the disease progression, generating microglial diversity and culminating with the so-called Disease-Associated Microglia (DAM) signature characterized, among others, by the upregulation of many genes identified by GWAS such as ApoE^7,8^.

Three ApoE alleles exist in the human population ε2, ε3 and ε4, ε4 being the strongest genetic risk factor of sporadic AD identified so far^9^. One of the proposed mechanisms by which ApoE could favor AD is through a direct interaction between Aß and ApoE. Studies have shown that ApoE can impact both Aß seeding, fribrillogenesis and clearance in an isoform dependent manner^10,11^, ApoE4 being more prone to facilitate seeding and to reduce Aß clearance^12,13^. ApoE functions in the CNS are nonetheless diverse and complex and likely contribute to AD through additional mechanisms. Recent data particularly revealed that both Trem2 and ApoE are critical regulators of microglial switch from homeostatic to neurodegenerative phenotype^14,15^ and genetic ablation of either gene results in a larger proportion of microglia in homeostatic state in mouse models of AD^15,16^.

Purinergic signaling is central to microglial biology in both healthy and pathological conditions^17^. Indeed, microglia express a large repertoire of purinergic receptors as well as different proteins involved in ATP release or degradation^17^. Microglial purinergic receptor expression is highly dependent on the state of microglia. In the homeostatic state, *P2ry12* gene is among the most expressed, while its expression is strongly down regulated in reactive states^18^. Conversely, P2X4 receptor, an ATP-gated channel, is not present in homeostatic microglia but its expression is induced upon activation^19^. As a consequence, reactive microglia loses or acquires functions associated with these two receptors.

In reactive microglia, P2X4 receptors have been linked to different functions and pathologies. In neuropathic pain models, *de novo* P2X4 expression in spinal cord microglia enhances local network excitability^20^. Similarly, following a *status epilepticus*, P2X4 hippocampal microglia likely contribute to microglial-evoked neuroinflammation and neuronal death^21^. Generally, pharmacological or genetic blockade of P2X4 receptors has beneficial effects in different acute CNS pathologies^22^.

However, potential involvement of P2X receptors in neurodegenerative diseases associated with inflammation is still poorly documented. Recently, the exploration of P2X7 in AD reveals detrimental functions, inducing chemokines release or T cells recruitment^23^. Whether P2X4 have detrimental of beneficial effects in slowly progressing neurodegenerative disease remains to be elucidated. Here, we investigated the potential role of P2X4 in AD. Our results show that in APP/PS1 mice, P2X4 is almost exclusively expressed in plaque associated microglia. Genetic deletion of *P2rx4* in APP/PS1 mice reverses cognitive declines and is associated with in a lower amount of soluble Aß1-42. In myeloid cells, P2X4 specifically interact with ApoE and triggers its degradation by regulating cathepsin B activity and *P2rx4* deletion results in higher amount of intracellular and secreted ApoE in both BMDM and microglia from APP/PS1 brain. Our results support that microglial P2X4 promotes lysosomal ApoE degradation, indirectly altering Aß peptide clearance, which in turn might promote synaptic dysfunctions and cognitive deficits.

## Results

### P2X4 is predominantly expressed in plaque-associated microglia

In most regions of the healthy brain, P2X4 receptors are expressed at low level except in a few regions such as the pyramidal cell layer of the hippocampus or the arcuate nucleus of the hypothalamus^24,25^, where a higher expression of the receptor was reported. Yet, in pathological conditions, both microglia and neurons might up regulate P2X4 expression^26^. Except few transcriptomic data, P2X4 expression in AD has not yet been observed^27^. We therefore analyzed whether P2X4 is upregulated in AD brain and determine in which regions and cell type. Using cortices slices of control and AD human patients, immunohistochemistry reveals a strong P2X4 immunostaining colocalized with Iba1, a specific marker of microglial cells, and amyloid plaques staining, while in control brain P2X4 staining was almost absent (**fig. 1A**). In the cortex of 12 months old APP/PS1 mice, P2X4 antibody reveals cluster of positive immunostaining, that was absent from WT mice. Co-staining of Iba1 shows a strong co-localization of P2X4 in microglia clustered around amyloid plaques deposit **(fig. 1B, C)**. No obvious P2X4 staining was found outside these patches **(fig. 1D)**. These results indicate that in AD brain, P2X4 is specifically up regulated in a subpopulation of reactive microglial cells, presumably in the so-called disease associated microglia^7,28^.

**Figure 1:**
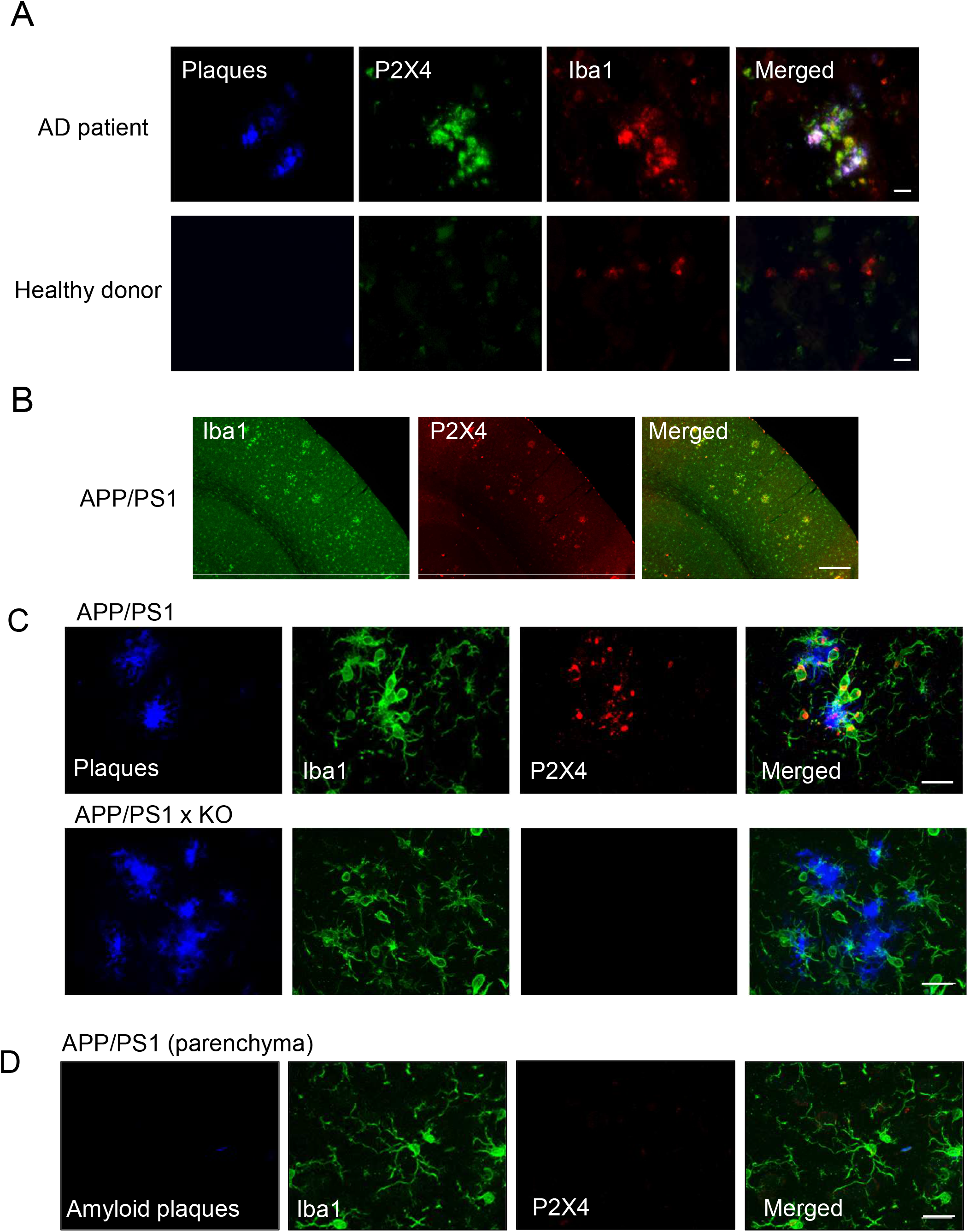
P2X4 is specifically expressed in plaque associated microglia in both human and mice AD brain. **(A)** Representative pictures of cortical brain slices from AD patients and healthy control labeled with AmyloGlo (blue, amyloid plaques), P2X4 (green) and Iba1 (red). P2X4 staining co-localizes with Iba1 in regions of dense amyloid plaque staining, supporting that microglia clustered around amyloid deposit specifically express P2X. In healthy control brain, P2X4 staining does not co-localizes with that of Iba1. Scale bar 20 μm. **(B)** Representative low magnification picture of immunofluorescence showing P2X4 (red) and Iba1 (green) immunostaining in the cortex of 12 months APP/PS1 mice. Both P2X4 and Iba1 staining co-localize in spot corresponding to amyloid plaques. Scale bar 200 μm. **(C)** High magnification of P2X4 (red) Iba1 (green) immunostaining at the vicinity of amyloid plaques (Amylo Glo staining, blue) in the cortex of 12 APP/PS1 mice *(top)* and APP/PS1xKO mice (*bottom*). Note the specific intracellular localization of P2X4 in microglia clustered around amyloid deposit. Scale bar 20 μm. **(D)** Representative immunofluorescence in APP/PS1 mice showing that parenchymal microglia (Iba1, green) do not express P2X4 (red) in region with no amyloid deposit (Amylo Glo staining, blue). Scale bar 20 μm.

### P2rx4 deletion reverses memory deficits in APP/PS1 mice

Alteration of cognitive performance is a hallmark of all mouse models of AD and particularly spatio-temporal disorientation is a major early sign of the disease. APP/PS1 mice display decline of cognitive performance in a variety of learning behavioral tests meant to assess spatial memory^7,28^. To address the role of P2X4 upregulation in APP/PS1 mice, we generated WT, P2X4^-/-^ (so called KO thereafter), APP/PS1 and APP/PS1xKO animals. First, we used the Hamlet test to assess topographical memory in the different groups of animals ^29,30^. 72 hours after the last training session, mice were water deprived (WD) for 15 hours. The probe test was performed by placing mice in the central agora for 10 min. Latency and number of errors to reach the drink house were analyzed. A second probe test was repeated the following day, animals being once again placed in the apparatus but in non-water deprived (NWD) condition. Both WD-WT and WD-KO mice showed a significant shorter latency to reach the drink house as compared to NWD condition, signing proper memory (**fig. 2, left panel,** WD-WT: 24.4 ± 7.3s; NWD-WT: 81.5 ± 1.7s; WD-KO: 13.9 ± 3.2s; NWD-KO: 40.9 ± 6s). As expected, WD-APP/PS1 mice did not shown difference as compared to NWD-APP/PS1 (WD-APP/PS1: 73.5 ± 24.5s and NWD-APP/PS1: 87.1 ± 19.5s), suggesting impaired memory. Remarkably, learning deficits observed in APP/PS1 mice were reverted in APP/PS1xKO mice (WD-APP/PS1xKO: 39 ± 9s and NWD APP/PS1xKO: 112.4 ± 22.1s) and memory was found similar to both WT and KO animals.

**Figure 2:**
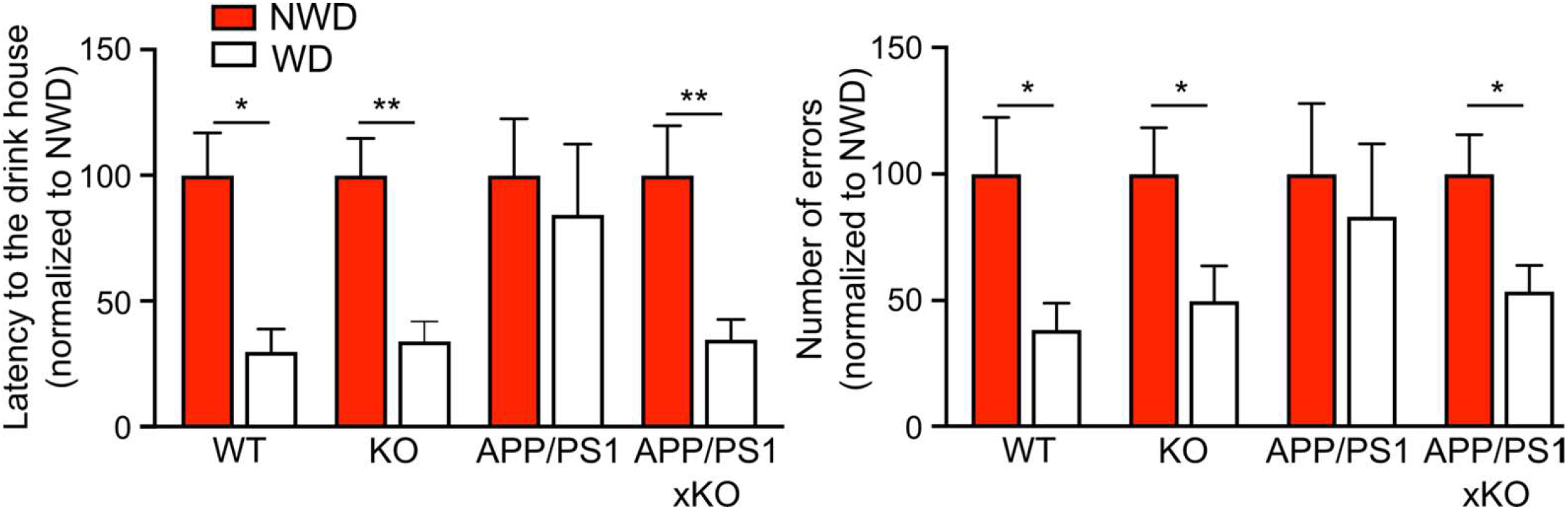
*p2x4* deletion reverses memory deficit in APP/PS1 mice. **(A)** *Left*, Latency to locate the drink house 15 h after water deprivation in the Hamlet test. WT and KO mice present reduced latency to the drink house, whereas no difference was observed between non-water deprived (NWD) and water-deprived (WD) conditions in APP/PS1 mice. APP/PS1xKO water deprived mice present significant reduction of the latency, indicating that mice have retained the location of the drink house. *Right*, Number of errors before entering the drink house. WT and KO mice present reduced number of errors, whereas no difference was observed between non-water and water-deprived condition in APP/PS1 mice. APP/PS1xKO deprived-water mice present significant reduction of the number of errors. N = 3 independent experiments, n = 8-11 mice per group. * p<0.05, ** p<0.01, Mann-Whitney test, WD *vs* NWD for each genotype.

Similar data were found when addressing the total number of errors. The later was reduced in both WD WT and KO mice as compared with NWD condition (WD-WT: 14.1 ± 3.9 *vs* NWD-WT: 36.8 ± 8.2; WD-KO: 11.1 ± 3.1 *vs* NWD-KO: 22.3 ± 4.1). This difference was absent in WD-APP/PS1 mice (WD: 33.5 ± 11.6 *vs* NWD: 40.3 ± 11.2), but readily observed in WD-APP/PS1x KO (WD: 23 ± 4.5 *vs* NWD: 43.7 ± 6.8) (**fig. 2, right panel**). Essentially similar findings were obtained in the Morris Water maze (Sup. fig. 1A, B, C). Locomotor activity of the different genotypes analyzed in the open field task indicated that both APP/PS1 and APP/PS1xKO mice presented a tendency to higher locomotion compared to WT and P2X4 KO, ruling out that the alteration observed in the Hamlet test could relate to mobility deficits (Sup. fig.1D). These results indicated that invalidation of P2X4 rescued memory deficits observed in APP/PS1 mice.

### P2X4 deletion reduces sAβ content

We next assessed whether the deletion of P2X4 could affect amyloid load in APP/PS1 mice. Number of plaques, their average size and the number of microglia associated with plaques were quantified in the cortex of 12 months APP/PS1 and APP/PS1xKO after AmyloGlo staining **(fig. 3A)**. As show in **figure 3B**, in APP/PS1 mice, P2X4 deletion did not altered density of plaques nor their average seize. The average number of microglia clustered around plaques was not changed between the two genotypes **(fig. 3C, D)**.

**Figure 3:**
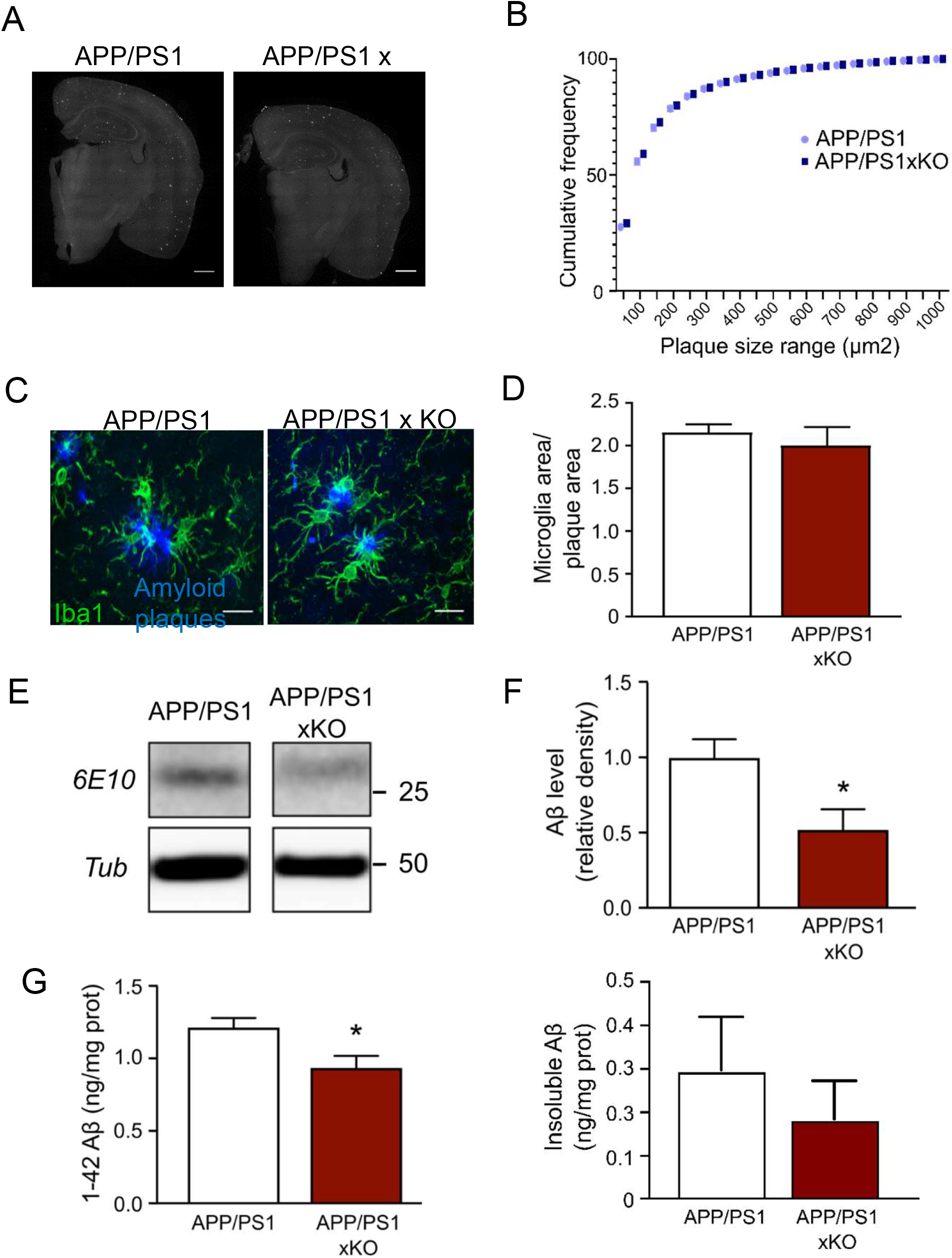
Deletion of *p2x4* does not affect amyloid plaques density but reduces soluble Aß species. **(A)** Representative images of Thioflavine T staining in APP/PS1 and APP/PS1xKO brain. Scale bar 700 μm. **(B)** Cumulative frequency of the range size of amyloid plaques. There is no obvious difference in the number of plaques not of their size between APP/PS1 and APP/PS1xKO; n = 11 mice per group. **(C and D)** Analysis of microglial clustering around amyloid plaque between in the cortex of APP/PS1 and APP/PS1xKO mice. (C) Representative image of microglia clustering around plaques. Amyloid plaques are stained in blue and Iba1 is in green. Scale bar 20 μm. **(D)** Quantification of the area covered by microglia surrounding amyloid plaques. The ratio of the surface occupied by microglia over the surface of the plaque is expressed for both APP/PS1 and APP/PS1xKO mice. n = 11 mice per group, unpaired t-test. **(E)** Representative Western blot of Aß peptide detected with the 6E10 antibody from cortex extracts from APP/PS1 and APP/PS1xKO mice. **(F)** Quantitative analysis of Western blots presented in (E). A significant decrease of the Aβ peptide amount is observed in APP/PS1xKO mice. n = 7 mice per group, * p<0.05, unpaired t-test. **(G)** ELISA quantification of soluble (right panel) and insoluble (left panel) Aβ1-42 peptide in the cortex of APP/PS1 and APP/PS1xKO mice. A significant decrease of the concentration of sAß is observed in APP/PS1xKO mice, compared to APP/PS1 mice. Insoluble Aß peptide is unchanged between the two genotypes. N = 2 independent experiments, n = 6-7 mice per group. * p<0.05, unpaired t test.

Finally, we analyzed if the level of the soluble Aß (sAß) peptide was affected by P2X4 deletion. Using Western blot analysis of cortical extracts, we found that the amount of sAβ was reduced in APP/PS1xKO compared to APP/PS1 mice (**fig. 3E, F,** 0.52 ± 0.1 *vs* 1 ± 0.1 respectively). Consistent with these findings, quantification of sAß_1-42_ peptide by ELISA confirmed the lower amount of soluble Aβ in the cortex of APP/PS1xKO compared to APP/PS1 (0.937 ± 0.083.6 ng/mg *vs* 1.216 ± 0.0665 ng/mg respectively, **fig. 3G**). All these results indicate that microglial P2X4 is associated with an increase of sAβ, which correlates with a decline of memory performances.

### P2X4 regulates cellular levels of ApoE

P2X4 receptors present a complex trafficking regulation with a prominent intracellular localization in the endo-lysosomal compartment^31^. We ask whether deletion of P2X4 could somehow alter endo-lysosomal functions therefore potentially regulating clearance of Aß peptide. To address this, we used an approach based on antibody-based affinity purification of native P2X4 receptors, followed by mass spectrometry to identify potential P2X4 partners involved in endo-lysosomal functions. To enrich specifically intracellular membrane compartments (i.e. endosome and lysosome) an ultracentrifugation step was performed after membrane protein solubilization (see methods section). Affinity purification was performed on bone marrow derived macrophages (BMDM) from both WT and P2X4^-/-^ mice. Among the different proteins interacting specifically with P2X4, ApoE was the only one with significative coverage found across two independent experiments **(sup. fig. 2)**. In BMDM, P2X4-ApoE interaction was further confirmed by immunoprecipitation using either P2X4 or ApoE antibodies (**fig 4A, B**).

**Figure 4:**
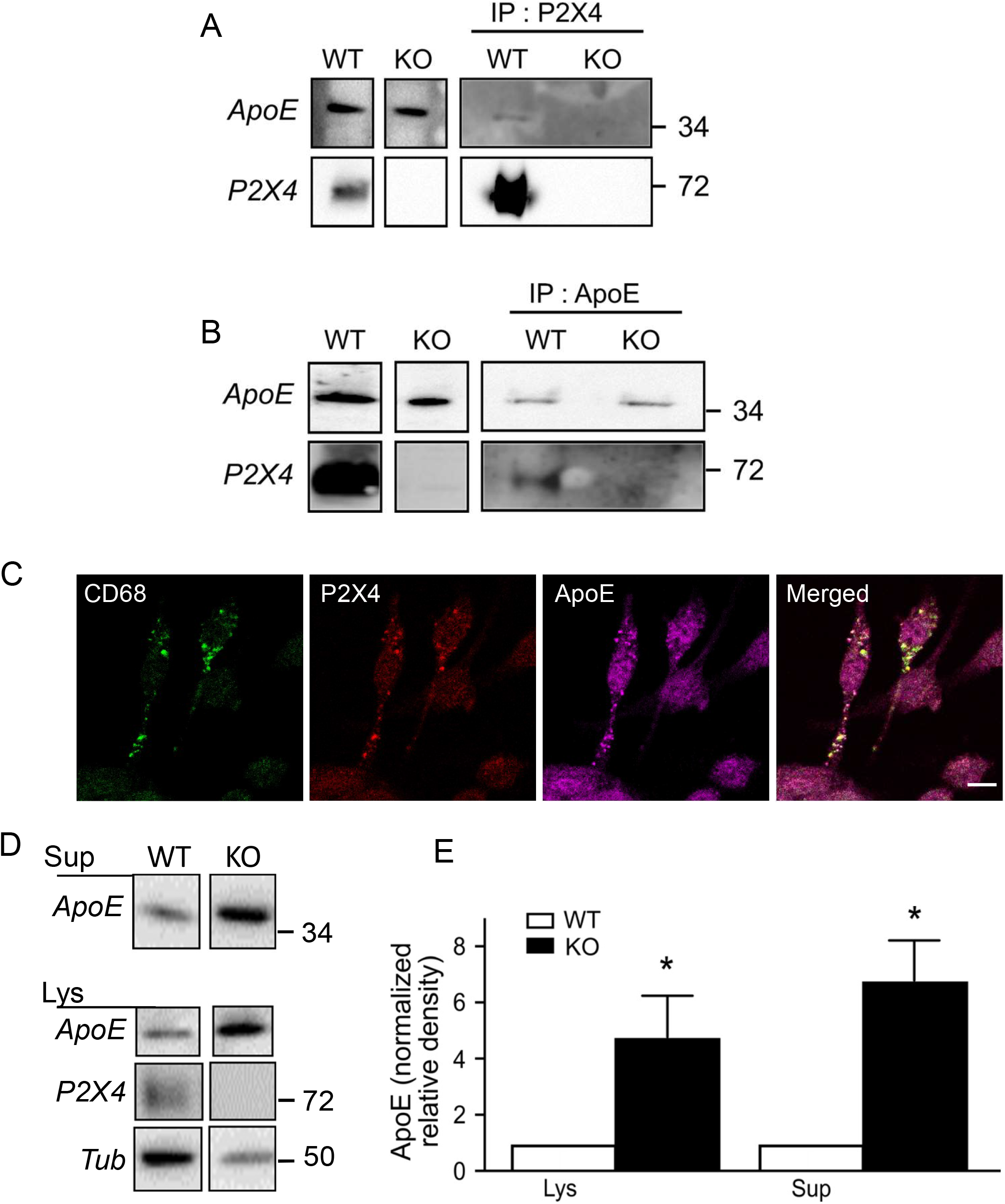
P2X4 interacts with ApoE in BMDM endo-lysosomal compartments and reduces its amount compared to P2X4-deficient cells. (**A, B**). Co-immunoprecipitation of P2X4 and ApoE. BMDM membrane extracts from WT and KO mice were immunoprecipitated (IP) with an anti-P2X4 antibody (A), or ApoE antibody (B). Immunoprecipitated proteins were separated by electrophoresis and immunoblotted with either anti-ApoE (top row) or anti-P2X4 (bottom row) antibodies. **(C)** Representative immunofluorescence image showing the co-localization of the lysosomal marker CD68 (green), P2X4 (red) and ApoE (purple) in BMDM cells. Scale bar 5 μm. **(D, E)** Western blot analysis of ApoE in BMDM culture supernatants (Sup) or cell lysates (Lys) from WT and KO mice. (D) Representative western blot of ApoE, **(E)** Quantification of western blot presented in D. A significant increase of ApoE is observed in both KO cultures supernatants and in cell lysates. Results were normalized to ApoE signal obtained for WT BMDM in each culture. n = 6 independent cultures, * p<0.05, unpaired t test.

Immunocytochemistry revealed that in BMDM, P2X4-ApoE interaction is localized in intracellular compartments, likely from the endo-lysosomal pathways. This was further demonstrated by the co-localization of P2X4 and ApoE with CD68, a specific marker of the endo-lysosomal pathway **(fig. 4C)**. The intracellular interaction between P2X4 and ApoE is consistent with the known presence of P2X4 in lysosomes. We thus investigated whether P2X4 could mediate the trafficking of ApoE to lysosome and contribute to its degradation. To that aim, we compared the expression of ApoE in BMDM from WT and P2X4-deficient mice by western blotting. Because ApoE is secreted by BMDM, amounts of ApoE was determined both in cell lysates and cell culture supernatants. As shown in **figure 4D, E**, BMDM from KO mice express much higher levels of ApoE in both cell lysates and supernatants. Compared to normalized WT values, levels of ApoE in KO BMDM were 4.7 ± 1.5 and 6.8 ± 1.4 fold higher in lysate and supernatant, respectively. Non-normalized data show the same results but with higher variability (**sup. fig. 3**). Essentially identical results were obtained in the lysate of primary culture of microglia from WT and KO mice (**sup. fig. 4**). RT-qPCR transcriptional analysis of ApoE mRNA from WT and KO BMDM did not reveal any difference **(sup fig. 5)** further supporting that the physical interaction between P2X4 and ApoE results in its degradation, likely through the lysosomal pathway. These effects of P2X4 on ApoE were reproduced in transfected COS-7 cells. Co-transfected cells show clear intracellular co-localization of P2X4 and ApoE (**fig. 5A**). As observed in BMDM, P2X4/ApoE co-transfected COS-7 cells presented significant lower amounts of ApoE in both lysates and supernatant compared to cells transfected with ApoE alone (**fig. 5B, C**).

**Figure 5:**
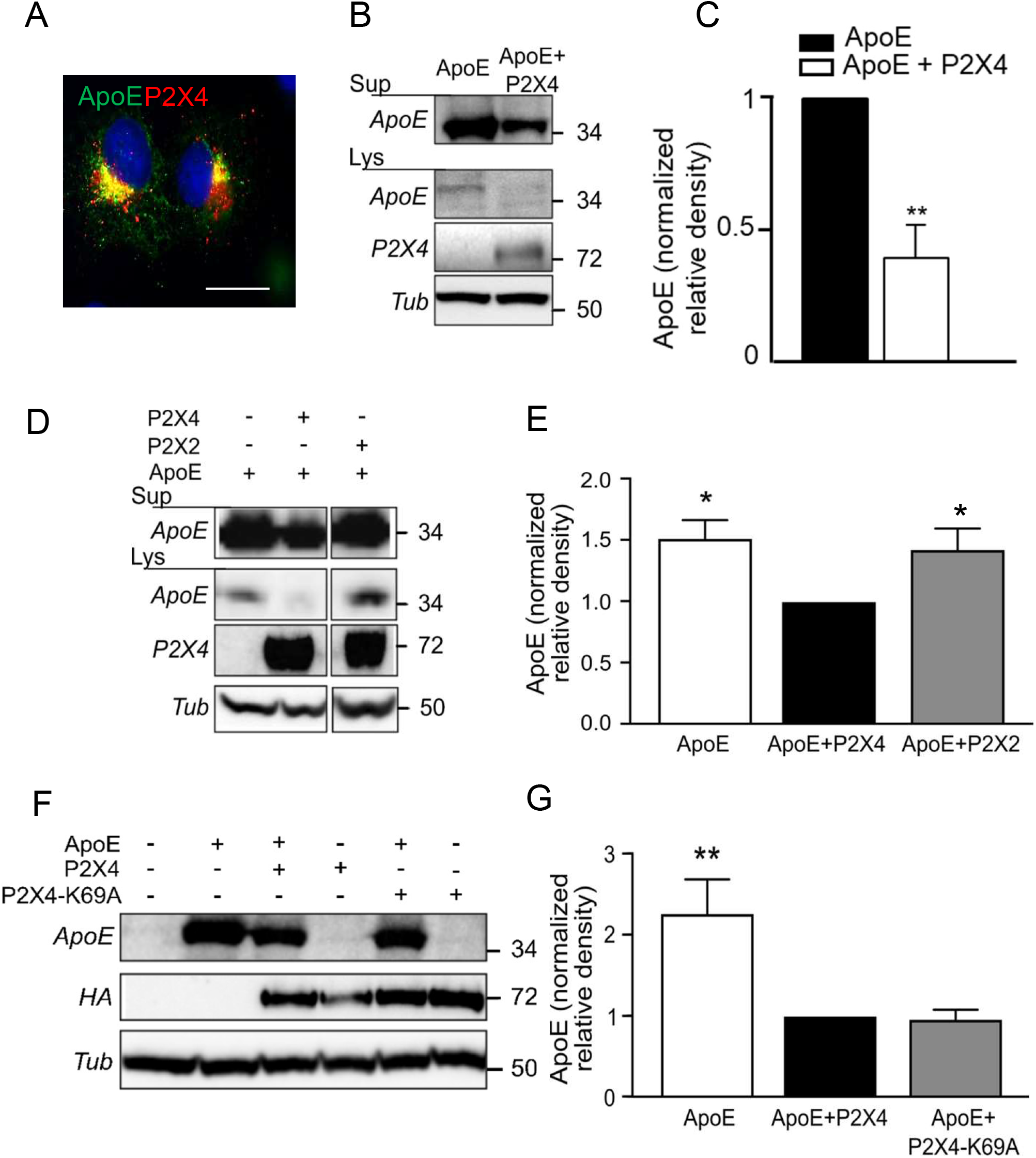
Characterization of the interaction between P2X4 and ApoE in recombinant system. **(A)** Representative immunofluorescence of ApoE (green) and P2X4 (red), and DAPI (blue) in co-transfected COS-7 cells. Both ApoE and P2X4 co-localize in intracellular compartments. Scale bar 10 μm. **(B, C)** Comparison of ApoE expression upon co-transfection with P2X4. COS-7 cells were transfected with ApoE alone or in combination with P2X4. Expression of ApoE was analyzed by Western blot in both cell culture supernatants and cell lysates (B). Quantitative analysis shows that in the presence of P2X4, amounts of ApoE is reduced in both culture supernatant (Sup) and cell lysates (Lys) (C). n = 3 independent experiments, ** p<0.01, One sample t-test compared to theoretical value of 1. **(D, E)** Comparison of ApoE expression upon co-transfection with P2X4 or P2X2. (D) Expression of ApoE was analyzed by Western blot in both cell culture supernatants and cell lysates (E) Quantitative analysis of ApoE in supernatant shows that co-expression with P2X4 reduces the expression of ApoE, whereas that of P2X2 has no effect. n = 6 independent experiments, * p<0.05, One sample Wilcoxon compared to theoretical value of 1. **(F, G)** P2X4 activity is not necessary to reduce ApoE levels. (F) Expression of ApoE was analyzed by Western blot in cell culture supernatants of cells transfected with ApoE alone or in combination of either P2X4 or P2X4-K69A, an ATP binding site dead mutant. (G) Quantitative analysis shows that both P2X4 and P2X4-K69A significantly reduces the ApoE levels to the same extend. n = 8 independent experiments, ** p<0.01, one sample *t*-test compared to theoretical value of 1.

We next examine whether others P2X receptor subunits could modulate ApoE levels. To address this question, ApoE contents were analyzed as above in COS-7 cells expressing ApoE alone or co-transfected with either P2X4 or P2X2. As shown in **fig. 5D, E,** as expected, co-expression of ApoE and P2X4 induced a reduction of both cellular and secreted ApoE as compared to cells expressing ApoE alone (0.95 ± 0.3 *vs* 1.34 ± 0.3) while co-expression of ApoE and P2X2 did not alter ApoE levels (1.18 ± 0.24 *vs* 1.34 ± 0.3, unnormalized values).

Finally, we ask whether P2X4-mediated ApoE downregulation was dependent on its channel activity. As above, levels of ApoE were measured in lysates and supernatants from COS-7 cells expressing ApoE alone or co-transfected with either P2X4 or P2X4-K69A, a mutant form of the receptor unable to bind ATP. As shown in **figure 5F, G**, levels of ApoE were the same in cells transfected with P2X4 or P2X4-K69A, but significantly lower than in cells expressing ApoE alone. These results suggest that P2X4 receptor activity is not necessary to drive down regulation of ApoE and are consistent with an intracellular mechanism.

### P2X4 induces cathepsin B activity

A potential explanation for the higher amount of ApoE in P2X4 deficient myeloid cells is that ApoE, through its interaction with P2X4, is trafficked to lysosomes where it is degraded. To test this hypothesis, we used the E64 compound, a known inhibitor of lysosomal proteases. WT and KO BMDM were incubated overnight with 10 μM E64, and levels of secreted and cellular ApoE were evaluated by western blotting. As shown on **fig. 6A and B**, in WT BMDM, E64 treatment strongly increased amounts of intracellular ApoE compared to untreated cells (0.44 ± 0.1 *vs* 0.23 ± 0.1, unnormalized values). In KO cells, E64 treatment had no effect on ApoE (0.92 ± 0.4 *vs* 0.81 ± 0.3). E64 is a broad-spectrum cysteine protease inhibitor which targets many different proteases either cytoplasmic or lysosomal. We therefore tested more specific inhibitors of cysteine proteases. We first evaluated the potential involvement of calpains, a family of mostly cytoplasmic proteases. Overnight pre-treatment of WT and P2X4-deficient BMDM with 10 μM calpain inhibitor III (CI-III), which targets calpains I and II, did not induced any significant change in ApoE amounts in either group (WT: 0.42 ± 0.2 *vs* CI-III: 0.43 ± 0.2; P2X4KO: 1.1 ± 0.5 *vs* CI-III: 1.57 ± 0.7) (**sup fig. 6A, B**). These results were confirmed using Suc-Leu-Leu-Val-Tyr-AMC, a fluorescent substrate of calpain. Incubation of BMDM with 100μM Suc-Leu-Leu-Val-Tyr-AMC showed no difference in fluorescence signal between genotypes (**sup fig. 6C**). These results indicate that calpains were not involved in ApoE degradation.

**Figure 6:**
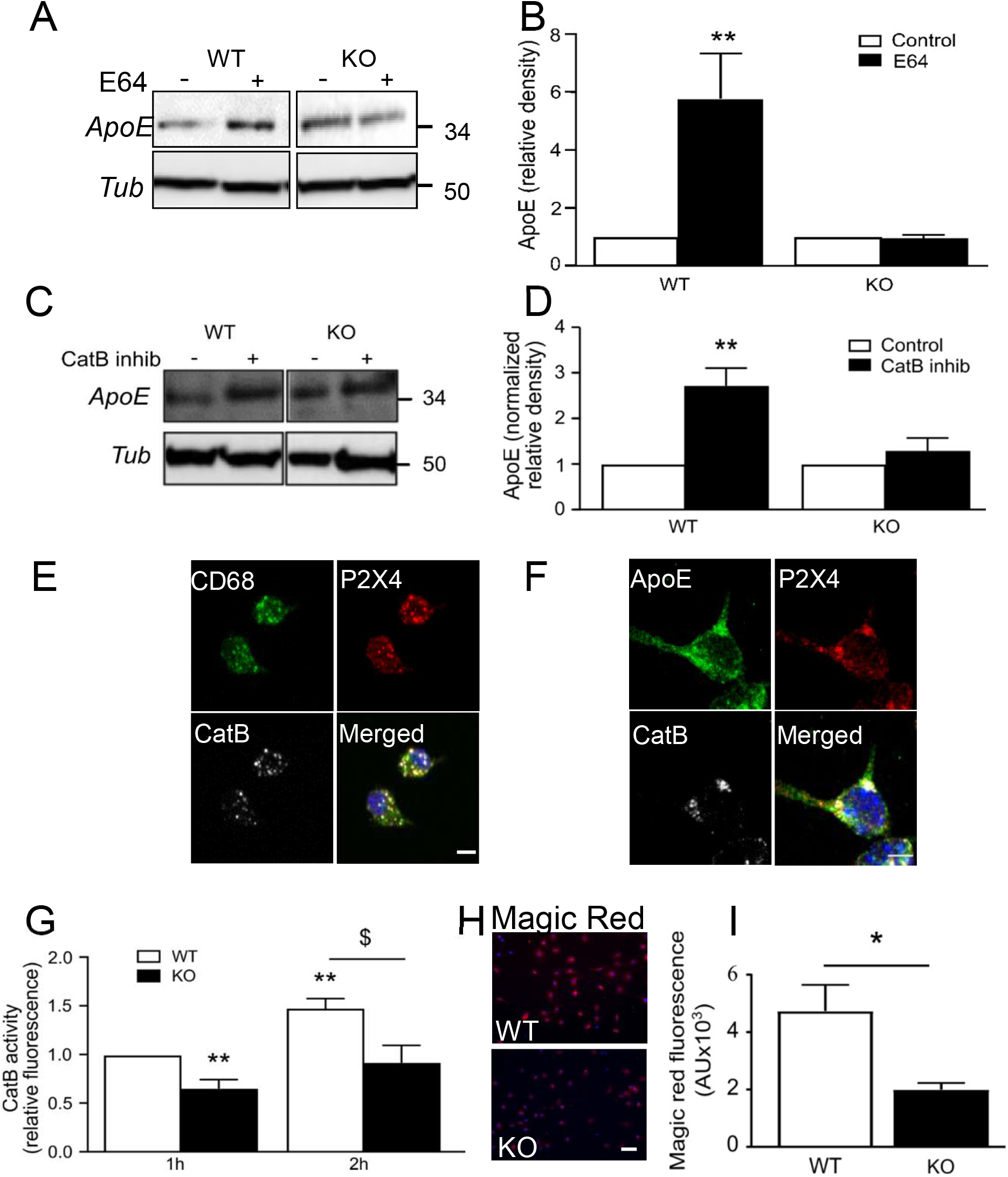
P2X4 regulates cathepsin B-dependent ApoE degradation. **(A, B)** Comparison of treatment with E64 a pharmacological inhibitor of the cysteine proteases, on ApoE expression in BMDM culture of WT and P2X4^-/-^ mice. (A) Representative western blot of ApoE in the supernatant of WT and P2X4^-/-^ BMDM after incubation with 10 μM E64. (B) Quantitative analysis of western blots shows that E64 induced a strong increase of ApoE in the supernatant of WT but not in P2X4^-/-^ BMDM. n= 5 independent experiments, ** p<0.01, One sample t-test compared to theoretical value of 1. **(C, D)** Comparison of treatment with 20 μM Z-Phe-Ala-FMK, a CatB inhibitor, on ApoE expression in BMDM culture of WT and P2X4^-/-^ mice. (C) Representative western blot of ApoE in the supernatant of WT and P2X4^-/-^ BMDM after incubation with Z-Phe-Ala-FMK. (D) Quantitative analysis shows that inhibition of CatB with Z-Phe-Ala-FMK induces a strong increase of ApoE in the supernatant of WT but not in P2X4^-/-^ BMDM. n = 6 experiments, ** p<0.01, One sample t-test compared to theoretical value of 1. **(E, F)** Co-localization in BMDM of P2X4, ApoE and CatB in CD68 positive compartments. (E) Representative picture of CD68 (green), P2X4 (red) and CatB (white) immunostaining in BMDM cells. Scale bar 5 μm. (F) Representative immunostaining of ApoE (green), P2X4 (red) and CatB (white) in BMDM cells. Scale bar 5 μm. **(G, H, I)** P2X4 regulates CatB activity in BMDM. (G) CatB activity was measured using the cell-permeable fluorogenic CatB substrate Z-RR-AMC. After incubation with 100 μM Z-RR-AMC, fluorescence was read at 1 h and 2 h. A significant increase of the signal is observed in WT macrophages between 1 h and 2 h, whereas the activity in KO cells remained unchanged. n = 8 experiments, ** p<0.01, One sample *t*-test compared to theoretical value of 1 WT(1 h) vs KO (1 h) and WT (1 h) vs WT(2 h), $ p<0,05 Kruskal-Wallis test WT(2 h) *vs* KO (2 h); KO (1 h) vs KO (2 h) is non-significant. (H) Representative microscopic image of cellular CatB activity in WT and P2X4^-/-^ BMDM using the Magic red cathepsin B kit. A strong signal is observed in WT BMDM as compared to P2X4^-/-^ cells. (I) Quantitative analysis of the magic Red fluorescence using ImageJ. * p<0.05, n = 3 independent cultures, unpaired *t*-test. Scale bar 30 μm.

We next tested whether cathepsin B (CatB), a cysteine protease highly expressed in lysosome, could be involved in ApoE degradation^32^. Pre-incubation of WT BMDM with 20 μM of CatB inhibitor overnight strongly enhanced amounts of ApoE (0.15 ± 0.1 *vs* 0.40 ± 0.2, unnormalized values) while having no effect on P2X4-deficient BMDM (0.44 ± 0.2 *vs* 0.46 ± 0.1, unnormalized values) (**fig. 6C, D**). No effect was observed using specific cathepsin L or cathepsin S inhibitors (data not shown). Triple immunostaining of P2X4, CD68 and CatB revealed a strong co-localization of the three proteins, indicating that P2X4 and CatB are both present in lysosome **(fig. 6E)**. A similar co-localization was observed for P2X4, CatB and ApoE (**fig. 6F**). Specificity of the CatB antibody was verified by immunostaining and western blot in CatB-deficient BMDM (**sup. fig. 7A, B**). We analyzed whether P2X4 deletion could alter the enzymatic activity of CatB. We first measured CatB activity using the specific CatB substrate ZZ-RR-AMC, which becomes fluorescent upon cleavage. WT and P2X4-deficient BMDM cells were incubated with 100 μM substrate for one and two hours and end point fluorescence was measured. As shown in **fig. 6G**, fluorescence was significantly higher in WT cells compared to P2X4-deficient BMDM, suggesting that CatB activity is reduced in these cells. These results were further confirmed with Magic Red assay, a cell-permeant CatB substrate whose fluorescence increases upon cleavage. **Fig. 6H, I** show that after incubation with Magic Red substrate, fluorescence was higher in WT compared to KO cells (47301 ± 9238 *vs* 19969 ± 2357 respectively, fluorescence arbitrary units). This lower CatB activity in P2X4-deleted cells was not due to an impaired expression of the enzyme since both WT and P2X4-deficient cells display similar amounts of CatB (**sup. fig. 7C**). Following our hypothesis that CatB controls the degradation of ApoE, we tested whether in BMDM from CatB-deficient mice would express higher level of ApoE. As shown in **sup. fig. 8**, a higher amount of ApoE in CatB-deficient BMDM compare to WT was observed.

### P2X4 regulates ApoE degradation in APP/PS1 mice

We next investigated whether in APP/PS1 mice, microglial P2X4 is also prone to regulate ApoE degradation as demonstrated *in vitro.* First, we analyzed localization of ApoE and P2X4 in 12-months old APP/PS1 mice. Triple cortical co-immunostaining of ApoE, P2X4 and Iba1 revealed that P2X4 co-localizes with ApoE in microglia that are clustered around plaques (**fig. 7A**), furthermore in microglia P2X4 co-localizes with CD68+ vesicles **(fig. 7B)**.

**Figure 7:**
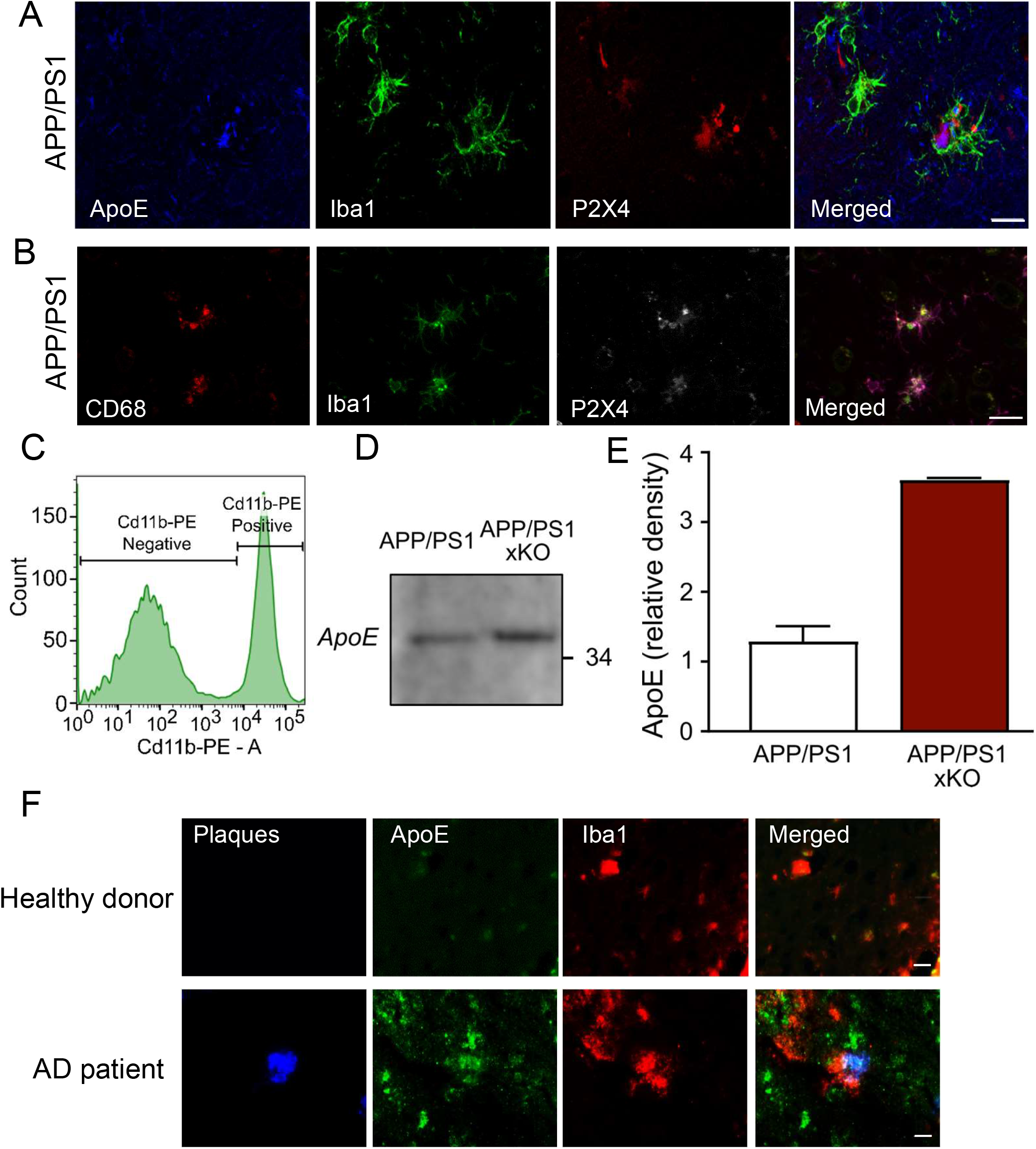
Increased ApoE in microglia from APP/PS1xKO mice and AD patients. **(A)** Immunofluorescence of ApoE (blue), Iba1 (green) and P2X4 (red) in APP/PS1 mice cortex. Scale bar 10 μm. **(B)** Immunofluorescence of CD68 (red), Iba1 (green), P2X4 (white) in APP/PS1 mice cortex. Scale bar 20 μm. **(C, D, E)** Analysis of ApoE expression in FACS sorted microglia from APP/PS1 and APP/PS1xKO mice. (C) Microglia were sorted based on CD11b-PE positive selection. (D) Representative western blot of ApoE from APP/PS1 and APP/PS1xKO FACS-sorted cortical microglia. **(E)** Quantitative analysis of signals presented in C shows an increase in ApoE in APP/PS1xKO mice relative to APP/PS1 mice. N = 2 independent experiments, n = 2 mice per group. **(F)** Representative pictures of cortical brain slices from healthy donor and AD patients labeled with AmyloGlo (blue, amyloid plaques), ApoE (green) and Iba1 (red) showing an increased expression of ApoE in human microglia clustered around amyloid deposit. Scale bar 20 μm.

We also quantified ApoE from FACS-purified CD11b+ microglia from APP/PS1 and APP/PS1xKO mice by western blot **(fig. 7C, D, E)**. Confirming previous *in vitro* findings, results show that the amount of ApoE was increased in purified microglia from APP/PS1xKO compared to APP/PS1 (0.36 ± 0.001 vs 0.13 ± 0.02, respectively). Finally, a clear co-localization of amyloid plaques, ApoE and Iba1 was also observed in brain of AD patient whereas in control brain, ApoE staining was almost slighter (**fig. 7F**), suggesting that in human AD brain, P2X4 could also regulate ApoE degradation. Altogether, our results indicate that microglial P2X4 receptors, independently of their pore activity, are involved in ApoE degradation by promoting the activity of lysosomal cathepsin B activity.

## Discussion

P2X receptors expression is up-regulated in reactive microglia associated with diverse neuropathological conditions such as neuropathic pain, *status epilepticus* or multiple sclerosis^22^. P2X4 activation in reactive microglia generally promotes deleterious effects such as hyperexcitability or inflammation^33^. Yet, in multiple sclerosis, P2X4 expression has beneficial effect by increasing myelin phagocytosis and favoring remyelination^34^. In this study, we investigated to what extend P2X4 contributes to microglial functions in Alzheimer’s disease.

By inactivating *p2x4* gene in APP/PS1 mice, we found that cognitive deficits associated with the APP/PS1 genotype were reversed to what is observed in wild type mice. This was associated with a reduction of soluble Aß peptide while there was no major difference in plaque load. Using a proteomic approach, we identified a specific interaction between ApoE and P2X4 in macrophages and further demonstrated that this interaction leads to a cathepsin B-dependent degradation of ApoE. In brain of APP/PS1 mice, P2X4 receptors are specifically expressed in so-called disease-associated microglia, a subpopulation of reactive microglia clustered at the vicinity of amyloid plaques, where they co-localize with ApoE. A similar pattern of expression was also found in post mortem human brain from AD patients. Altogether, our results support a role for P2X4 to promote microglial ApoE degradation which leads to sAß accumulation in brain parenchyma and contributes to memory deficit in APP/PS1 mice.

### *p2rx4* deletion reverses cognitive performance decline in APP/PS1 mice

We used both spatial memory as measured in water-maze learning and topographic memory in the Hamlet test, a recent behavioral device previously shown to measure spatio-temporal disrientation in mice^29^. In this test, APP/PS1 mice show a strong deficit to find the drinking house, a deficit which is no longer present in APP/PS1xP2X4^-/-^. *p2rx4* deletion has been linked to alteration of synaptic plasticity, which could result in spatial memory deficit^35^. However, our data supported that, in both the Hamlet test and water maze test, P2X4 deficient mice do not show learning impairment nor retention deficit. If some cognitive deficits have been reported in P2X4^-/-^ mice, these deficits relate to socio-communicative and sensorimotor impairments rather than to memory performance^36^.

In physiological conditions, P2X4 is expressed at low level in different neuronal populations throughout the brain, but absent from microglial cells^24^. In pathological conditions, P2X4 is expressed *de novo* in reactive microglia where it contributes to BDNF release, network excitability and inflammatory response^25,37^. In 9 months old APP/PS1 mice our data clearly show a strong expression of P2X4 in reactive plaque-associated microglia (PAM), while in aged match control mice, the expression of the receptor is barely detectable in brain parenchyma. Increased expression of P2X4 in PAM is supported by recent transcriptomic data which show that in laser captured plaque associated microglia, *p2rx4* is expressed more than 4-fold compared to microglia from the parenchyma, away from any visible plaque (Hemonnot et al., submitted). Although we cannot exclude a contribution of neuronal P2X4 receptor, our observations that (i) the receptor is highly expressed in PAM microglia and, (ii) its deletion reverse cognitive deficits of APP/PS1 mice but not in WT mice strongly support that microglial P2X4 receptor directly contributes to topographic and spatial memory alterations in AD mice.

Our results show that P2X4 deletion in APP/PS1 mice does not significantly change the number of amyloid plaques, nor the number of microglia in the parenchyma. Yet in APP/PS1xKO the amount of hippocampal soluble Aβ was significantly reduced compared to APP/PS1 mice in both western blot and ELISA experiments, while insoluble fraction was not different between the two genotypes (not shown). There are compelling evidence that toxic soluble low molecular-weight amyloid-beta ß oligomers directly induce synaptic deficit leading to a reduction of learning capacities^38^. The reduction of the amount of sAß observed in APP/PS1xKO could explain their better cognitive performances compared to APP/PS1 mice. Western blot analysis of synaptic proteins in APP/PS1 confirm a reduction of both NR1A and PSD95 relative to WT mice. However, a similar reduction was also observed for P2X4 deficient mice and APP/PS1xKO (not shown). One interpretation for this discrepancy is that the reduced level of sAß observed in APP/PS1xKO may be sufficient to restore synaptic efficiency independently of synaptic structural changes, e.g. by directly regulating neuronal excitability alterations associated with Aß peptide deposit^39^.

### P2X4 interact with ApoE and mediates its degradation

P2X4 receptors trafficking is tightly regulated and is mainly located in the endosomal/lysosomal network, which structural dysregulation in AD could promote abnormal APP processing^40^. Using a proteomic strategy based on intracellular organelle enrichment^41,42^, we identified ApoE as a specific P2X4 interacting protein in myeloid cells. Both proteins colocalize in intracellular CD68 positive compartments, likely endo-lysosome. Deletion of *p2x4* results in higher amounts of intracellular and secreted ApoE supporting that P2X4 drives CatB-dependent ApoE degradation. This interpretation is based on the observation that i) in P2X4 deficient macrophages and microglia, ApoE levels are increased compared to WT, independently of any transcriptional alteration, ii) inhibition of CatB enhances extracellular amounts of ApoE, an effect that is not observed in P2X4 deficient cells, iii) CatB activity is reduced in P2X4-deficient cells. Surprisingly, in recombinant system, introducing a binding site blocking mutation in P2X4 does not alter ApoE degradation, suggesting that P2X4 activity is not required. However, we cannot rule out an activity-dependent mechanism. Indeed, in the endo-lysosomal pathway ATP-binding region of P2X4 faces the organelle’s lumen and high ATP concentrations and acidic pH reduces P2X4 affinity for ATP^40^, it is conceivable that millimolar concentration of ATP in lysosome can trigger channel activity. Alternatively, alkalinization of lysosome may lead to P2X4 activation as previously shown, however such an activation would also lead to decrease CatB activity, which *in fine* would reduce ApoE degradation. Finally, using the fluorescent Lysosensor™, we did not observed any variation of intra-lysosomal pH in P2X4-deficient cells, further supporting that regulation of CatB activity by P2X4 is independent of pH variation. Yet, in the context of AD where dysregulation of lysosomal pH is well documented^43^, we cannot exclude that an activity of lysosomal P2X4 due to defective lysosomal acidification, could contribute somehow to ApoE degradation.

A key feature of P2X4 is its *de novo* expression in reactive microglia in diverse pathological conditions. Here, we show that in APP/PS1 mice, P2X4 is almost exclusively expressed in plaque associated microglia, but not in parenchymal microglial away from plaques nor in neurons. It is likely that P2X4 belongs to the so-called Disease Associated Microglia (DAM)^28^, a specific microglial population that is characterized by the specific expression of subset of genes, including several known AD risk factor such as Trem2 and ApoE. Indeed our data show a strong co-localization of P2X4 and ApoE in plaque associated microglia, in both mice and human AD patients, while in a recent study using mass spectrometry to identify deregulated proteins in microglia, an increase of P2X4 was observed in two mouse models of AD (APP/PS1 and APP-NL-G-F)^44^. While a clear *de novo* P2X4 protein expression is observed in reactive microglia, *p2rx4* gene has not been identified as deregulated in the various high throughput single cell genomics studies of reactive microglia. We made a similar observation in our previous transcriptomic analysis of reactive microglia in a model of sepsis^45^. This discrepancy between protein and RNA might be due to a translational regulation of P2X4 mRNA as previously suggested^46^.

### Role for microglial P2X4 /ApoE in AD

Our results show that in AD mouse brain, P2X4 is specifically expressed in microglia clustered around plaque that also express ApoE. P2X4 deficiency results in a significantly higher level of microglial ApoE, and presumably of its secreted form. P2X4 deficiency also leads to lower the amount of sAß. A great wealth of studies support a direct role of ApoE on sAß clearance^47^. It is surprising to observe that elevated levels of microglial ApoE correlates with reduced sAß and better cognitive performance in APP/PS1xP2X4KO mice. Indeed, mouse ApoE is thought to be amylogenic since global knock-out of *Apoe* results is dramatic reduction of Aß peptide deposition, even though deletion of *Apoe* show opposite effect on Aß deposition depending on the type of APP overexpressing mice model used^48–50^. However, a recent study demonstrated that microglial-specific inactivation of ApoE, beside a slight increase of average plaque size, has only limited repercussion of amyloid burden in the 5xFAD model^51^. Yet, microglial *apoe* deletion results in an age-dependent reduction of the synaptic markers PSD95 and synaptophysin, regardless of 5xFAD genotype. Of note, global *apoe* deletion promotes neuritic dystrophy^52^. Our results show that deletion of P2X4 increases microglial ApoE, reduces sAß and reverses cognitive deficits, further supporting a minimal role of microglial ApoE in amyloid plaque formation but a potential protective function toward synapses. Although our results do not allow to directly link increased levels of microglial ApoE in P2X4 deficient mice to the reduction of sAß or to the attenuation of the cognitive dysfunctions, the upregulation of P2X4 receptors in plaque associated microglia is likely involved in the development of AD behavioral deficits, probably by promoting ApoE degradation.

In human post mortem AD brain, we observed a similar distribution of P2X4, microglia and ApoE around amyloid plaques than in APP/PS1 mouse brain. This suggest that P2X4 could play similar functions in the human pathology, although AD mice models only partially recapitulate the human disease and mouse and human ApoE differently contribute to the disease^47^. Further experiments will be necessary to investigate this possibility.

Our data further support an important contribution of microglial P2X4 receptor to brain pathologies such as neuropathic pain, epilepsy or stroke. They also underline a potential protective function of microglial ApoE toward neurons cognitive performances.

## Methods

### Animals

Mice carrying a targeted null mutation of the P2RX4 gene were described elsewhere^35^. Briefly, a *E. Coli* ß-galactosidase (LacZ)-neomycin cassette was inserted in place of the first coding exon of the P2RX4 gene. In the resulting allele, the P2RX4 promoter drives ß-galactosidase expression. Chimeric mice were generated and crossed with C57BL/6 females to generate heterozygotes, which were then intercrossed to give rise to overtly healthy offspring in the expected Mendelian ratio. In the present study, mice were backcrossed for at least 20 generations and then maintained as separate P2RX4 knockout (P2RX4^-/-^) and wild-type (P2RX4^+/+^) lines. All experiments followed European Union (Council directive 86/609EEC) and institutional guidelines for laboratory animal care and use. Institutional license for hosting animals was approved by the French Ministry of Agriculture (No. D34-172-13). The Tg(APPswe,PSEN1dE9)85Dbo mice^53^ (APP/PS1) were obtained from the Jackson Laboratory (JAX stock #034832) and bred as heterozygotes to C56 Bl6/J or P2X4^-/-^ mice. All experiments using APP/PS1 and APP/PS1xP2X4^-/-^ mice were carried out at 12 months of age.

### Behavioral experiments

#### Hamlet test

The Hamlet test was performed as previously described^29,30^. Briefly, the device consisted of a 1.6 m diameter apparatus with an agora in the center and five corridors expanding toward different compartments, called houses. Each house has a different interest: drink, eat, run, hide or interact with a stranger mouse. Mice were trained in group and were allowed to go freely in the apparatus for 4 h per day during 12 days. Probe tests were performed 72 h and 96 h after the last training day, in water deprived or non-water deprived conditions, respectively. For water deprived condition, water bottles were removed from mice housing cages 15 h before the test. Mice were placed in the agora for 10 min and exploratory behaviors were video-tracked and analyzed with the Viewpoint software as latency time and number of errors to go to the drink house.

#### Morris water maze

The Morris water maze test was performed in a 1.4 m diameter (40 cm height) circular tank with extra maze cues. Tank was filled with 22°C water containing non-toxic lime carbonate to make it opaque. A 10 cm diameter circular platform was immerged under water, thus not visible to mice. Mice were trained three times a day for six consecutive days. They were allowed a free 90 s swim in order to find the platform. If by that time, mice did not find the platform, they were gently place on it and stayed there for 20 s. Probe test were performed 48 h after the last training day. The platform was removed and mice swam for 60 s. A video camera recorded the probe test and analysis was performed using the Viewpoint software.

#### Locomotor activity

Mice were place in a square open field box for 10 min. Viewpoint software tracked animals and calculated the distance travelled.

### Tissue preparation

Mice were euthanized with Euthasol (300 mg/kg) and perfused with PBS. Brains were either collected and fixed in 4% PFA at 4°C overnight or stored at - 80°C.

### Cell culture and transfection

COS-7 cells were cultured in Dulbecco’s Modified Eagle (DMEM) + glutamaX buffer supplemented with 10% fetal bovine serum (FBS) and 1% Penicillin-Streptomycin, and kept at 37°C with 5% CO_2_. Before transfection, cells were plated at 70% confluence in 6-well plates. Transfection was carried out using Lipofectamine 2000 (ThermoFisher), with the following DNA amount: 150 ng ApoE, 100 ng *p2rx4*, 100 ng *p2rx4K69A* and 100 ng *p2rx2*. Medium was changed for HBSS (Gibco, 14025092) 48 h after transfection and supernatants and cell extracts were collected the next day as described below.

### BMDM culture

BMDM were obtained from mice femur and tibia bone marrow and cultured in 30% L929 cell media and 70% DMEM + glutamaX buffer, supplemented with 10% fetal bovine serum (FBS) and 1% Penicillin-Streptomycin. Cells were mechanically dissociated and plated and medium was changed every 3 days. BMDM from cathepsin B deficient mice^54^ were kindly provided by Dr. Bénédicte Manoury (Hôpital Necker Enfants Malades, Paris).

### Pharmacological treatment

Macrophages were plated in a 12-well plate at 10^6^ cells per well and treated with 10 μM E64 (Tocris, 5208) or 20 μM Z-Phe-Ala-FMK cathepsin B inhibitor (Santa Cruz, sc3131) in HBSS overnight.

### Western blot

For BMDM culture cells, supernatants were collected in Amicon column (Millipore, UFC5010BK) and centrifuged at 14000 *g* for 30 min at 4°C. Column fraction were then collected and constituted the supernatant fraction of our cells. Cells were homogenized in lysis buffer (100 mM NaCl, 20 mM HEPES, 5 mM EDTA, 1% IGEPAL containing protease inhibitors). For cortex samples, dissected cortices were mechanically homogenized in 1% Triton lysis buffer (100 mM NaCl, 20 mM HEPES, 5 mM EDTA, 1% Triton X100 containing protease inhibitors) before homogenization on a wheel at 4°C for 1 h. Protein extracts were then centrifuged at 15000 *g* at 4°C for 10 min. After measuring protein concentration using Bradford technique, LDS sample buffer and 10% β-mercapto-ethanol were added. Proteins were then separated by reducing 4-12%, SDS-PAGE and transferred to a nitrocellulose membrane. The membrane was blocked with PBS with 0.1% Tween 20 (PBST) containing 5% non-fat dry milk overnight at 4°C. The membrane was then incubated overnight at 4°C with the indicated antibodies. After three washes in PBST, the membrane was then treated for 45 min at room temperature with the appropriated HRP-conjugated secondary antibody: Proteins were visualized using an ECL+ detection kit (Amersham) and imaged using a Chemidoc Touch Imaging system (Biorad). Densitometry was analyzed using the ImageLab software.

### Human tissue

Frozen brain samples from human tissue were obtained by the IHU-A-ICM-Neuro-CEB brain bank (Hôpital de la Pitié-Salpétrière, Paris). For immunostaining, cortex slice arrived frozen and mounted on microscope slides.

### Aβ ELISA

Cortices were homogenized in Tris buffer (Tris 1M, pH 7.6-SDS 2%) containing protease and phosphatase inhibitors and 1mM AEBSF. Homogenates were then sonicated at 40 mV for 10 s and centrifuged at 13000 *g* for 30 min at 4°C. Supernatants were then collected and constituted the soluble fraction of the sample. Quantification of soluble Aβ peptide was performed using an ELISA kit (Thermofisher, KHB3441).

### Immunostaining

Tissues were cut using a vibratome into 40 μm sections, rinsed with PBS and blocked with 10% goat serum diluted in a 0.1% Triton X100 solution for 2 h. Appropriate primary antibody was added overnight at 4°C. After rinsing, slices were incubated for 2 h with corresponding secondary antibody. Antibodies were diluted in PBS with 0.1% TritonX100. Amylo Glo (Biosensis, TR300-AG) was used for amyloid plaque staining according to the provided instructions. Briefly, before immunostaining, brain section were transferred in a 70% ethanol solution, rinsed with distilled water and incubated with Amylo Glo for 10 min. Before mounting, sections were incubated with 1x True Black^®^ (Biotium) to quench lipofusin autofluorescence. After rinsing, sections were mounted with Fluorescent Mounting medium (Dako) and observed on an AxioImager Z1 apotome (Zeiss).

BMDM cells were fixed in 1% PFA, washed and incubated with 10% goat or donkey serum in PBS containing 0,1% Triton X-100 for 30 min. Cells were then incubated with primary antibodies for 2 h, washed and incubated with secondary antibodies for 1 h before mounting.

Human tissues were fixed with 4% PFA and incubated with 10% goat or donkey serum in PBS containing 0.1% Triton X-100. Primary antibodies were directly put on slides in PBS containing 0.1% Triton X-100 overnight. After washing three times in PBS, tissues were incubated with the secondary antibodies for 2 h.

### Amyloid plaque quantification

Amyloid plaques size was quantified using Thioflavine T staining. Brain section were stained with 100 μg/mL Thioflavine T (Sigma T3516) for 15 min, rinsed with ethanol 70% for 5 min once and with PBS three times. Brain sections were mounted and images were acquired using a Zeiss AxioImager Z1 microscope. Plaque size was quantified using the threshold function in ImageJ. Then frequency was calculated using the frequency function in Excel. For each animal, 5 brains sections were analyzed.

### Microglia area quantification

Brain sections were stained with AmyloGlo® and Iba1 antibody in order to stained amyloid plaques and microglia. For quantification, fields containing plaques were randomly chosen; six fields per section, five sections per animals were acquired using a Zeiss AxioImager Z1microscope. For each amyloid plaque, the field of interest analyzed is defined by a perimeter that is proportional to the plaque size: the perimeter is calculated with a radius equal to four time the radius of the amyloid plaque. The Iba1 area is quantified in the zone using the threshold function in ImageJ.

### Primary and secondary antibodies

The following antibodies were used: goat anti-ApoE (1:1000, Millipore AB947), goat anti-Cathepsin B (1:1000, R&D systems AF965), rabbit anti-P2X4 (1:200, Sigma, HPA039494), donkey anti-CD68 (1:300, Biorad, MCA1957A488T), mouse anti-6E10 (Biolegend SIG-39320-0200), mouse anti-tubulin (Sigma, T9026), rabbit anti-HA (Invitrogen 715500), rabbit anti-Iba1 (1:2000, Wako MNK4428), rat anti-P2X4 (1:200, kindly provided by Dr. Nolte^55^), donkey anti-rat-A594 (1:500 Jackson Immunoresearch 712-586-150), donkey anti-goat-A488 (1:2000, Molecular probes A11055), donkey anti-rabbit-A557 (1:2000, R&D systems NL004), goat anti-rabbit-A488 (1:2000, Molecular probes A11034), horse anti-mouse-HRP (1:2000, Cell signaling, 7076S), goat anti-rabbit-HRP (1:2000, Jackson, 111 −035-144), donkey anti-goat-HRP (1:2000, Jackson, 705-035-003).

### Cathepsin B fluorescent activity

BMDM were plated in a 96-well plate at 10^5^ cells per well and incubated with 100μM of the cathepsin B substrate Z-RR-AMC (Enzo, BML-P137) for 1h and 2h before reading fluorescence (ex 365, em 440) on a plate reader (Flexstation 3, Molecular Devices,). For cathepsin B activity assessment by microscopy, macrophages were plated in a 24-well plate containing cover slips and incubated with the Magic Red cathepsin B substrate (1/250) (Bio-Rad, ICT938) for 2h. Cells were then fixed and mean fluorescence intensity in cells was quantified using the ImageJ software.

### Membrane solubilization

Plasma membrane-enriched protein fractions were prepared from freshly isolated mouse BMDM. BMDM were detached by cell scraping, counted, pelleted and homogenized in a solubilization buffer (0.32M sucrose, 10mM Hepes, 2mM EDTA and complete protease inhibitor cocktail, pH 7.4) with 150 strokes of a Potter-Elvehjem homogenizer (Dominique Dutcher). The homogenates were centrifuged 20 min, at 1000 g at 4°C to eliminate the debris and the supernatant was centrifuged 1 h at 70000 *g* at 4°C. The supernatant was discarded and the resulting pellet was solubilized in a set of detergent buffers of variable stringency (Complexiolyte (Logopharm), CL48 was chosen for further experiments due to its physiological stringency and after analysis of its solubilization efficiency) supplemented with protease inhibitors (Roche) for 2 h at 4°C. Insolubilized material (pellet; particles > 336 S) was removed by centrifugation (30000 *g*, 18 min, 4°C) leaving micelles with an estimated size (diameter) of up to 75 nm in the supernatant.

### Purification of P2X4 receptor complex by immunoprecipitation and analysis by mass spectrometry

Freshly prepared solubilized proteins were incubated o/n at 4°C with affinity-purified rabbit anti-P2X4 antibodies (Alomone) cross-linked to magnetic beads (Dynabeads, Invitrogen). The flow through was discarded and the beads washed 5 times with wash buffer (CL48 diluted ¼ in PBS and supplemented with complete protease inhibitor cocktail (Roche)) and sample buffer (Invitrogen) was added to separate the protein complexes from the beads. Eluates were shortly run on SDS/PAGE gels, Coomassie blue stained and sliced according to molecular mass. Further treatments and tandem mass spectrometry was performed at the Harvard Medical School Taplin Biological Mass spectrometry facility, Boston, MA, USA.

### Co-immunoprecipitation

Experiments with BMDM were carried out with membrane-enriched protein fractions (see protocol above). COS-7 cells were homogenized in lysis buffer (100 mM NaCl, 20 mM Hepes, 5 mM EDTA, 1% NP-40 and complete protease inhibitor cocktail pH 7.4) 48 h after transfection. Lysates were clarified by centrifugation. Protein concentration of the lysates was determined using a protein assay kit (Bio-Rad) and were incubated on a rotating wheel with either specific antibodies crosslinked to magnetic beads (Dynabeads, Invitrogen) (rabbit anti-P2X4 (Alomone Labs), rat anti-P2X4 (kind gift of F. Nolte (Universitätsklinikum Hamburg-Eppendorf)), goat anti-ApoE (Millipore)) or anti-HA beads (Santa-Cruz Biotechnology) for 4°C, o/n. After five washes in lysis buffer, bound proteins were eluted with sample buffer (Invitrogen).

### Cytometry

Mice were perfused with PBS and cortices were collected and dissociated using the Neural Tissue Dissociation Kit P (Miltenyi, 130-092-628) combined with the gentleMACS Octo Dissociator with heaters, as indicated by the supplier. After dissociation, myelin was removed using the Debris Removal Solution (Miltenyi, 130-109-398). Cells were then incubated with Fc bloc (1:100, BD Pharmigen 553142) for 10 min on ice and stained with Cd11b-PE (BD Pharmigen, 557397) for 30 min on ice. Cells were first discriminated with size and granularity. Microglia were then sorted using a laser with a 561 nm excitation wavelength and a 582nm filter. Afterward, sorted microglia were homogenized as described above and used in western blot experiments.

### Quantitative PCR analysis

Total RNA from macrophage cultures was extracted with RNeasy Mini Kit (Qiagen). A measure of 2 μg of total RNA were reverse transcribed using random hexamers and SuperScript III First-Strand synthesis System (Invitrogen) according to the manufacturer’s instructions. Realtime PCR was performed by using SYBR Green dye detection according to the manufacturer’s instruction (SYBR Green PCR Master Mix, Roche) on a LightCycler480 system. PCR reactions were performed in a 10 ml volume containing 2.5 ml of diluted RT product, 1 ml of forward and reverse primers and 5 ml of PCR master mix. Negative controls using non-reverse-transcribed RNAwere performed simultaneously. For each reaction, Cq was determined using the 2nd Derivative Max tool of LightCycler480 software. The relative ratios of specifically amplified cDNAs were calculated using the DCq method (Pfaffl, 2001). RNAs from three independent cultures were used. All experimental conditions were processed at the same time.

### Measurement of lysosomal pH

BMDM were plated on 96 well plates and incubated with Lysosensor™ (Invitrogen, L7545) at 3 μM for 3 min at 37°C. Cells were then rinsed with PBS twice. A calibration curve of the intensity fluorescence as a function of pH was made. In order to do so, after incubation with Lysosensor™, cells were incubated with determined pH solution for 10 min at 37°C. Fluorescence was determined using a plate reader Spark (Tecan) using 340 nm and 380 nm excitation wavelength. Then ratio between fluorescence intensity resulting from the 340 nm and 380 nm excitation were calculated and pH was determined using the calibration curve.

### Statistics

Statistics tests were performed using the GraphPad Prism9 software. After checking that all parametric assumptions were met, data were analyzed using Student’s or ANOVA test. When the assumptions were not met, Wilcoxon signed-rank or Kruskal-Wallis test were used. For pharmacological treatment, data were paired. For each graph, mean ± SEM are indicated.

## Acknowledgements

This work was supported by the Institut National de la Santé et de la Recherche Médicale (INSERM), the Centre National de la Recherche Scientifique (CNRS), “La fondation NRJ-Institut de France”. JH was supported by LabEx ICST Grant ANR-11-LABX-0015. We thank Dr. Hélène Hirbec for her help with qPCR experiment. Experiments were performed with the help of the following Montpellier Biocampus core facilities: Réseau des Animaleries de Montpellier (RAM) (iexplore facility), the imaging facility MRI, member of the national infrastructure France-BioImaging infrastructure is supported by the French National Research Agency (ANR-10-INBS-04, «Investments for the future»). We also thank the CECEMA animal facility (University of Montpellier, France).

## Authors contribution

F.R. and L.U. conceived the study. J.H., E.G.P, L.U., F.R designed experiments. J.H., E.G.P, N.L., B.M., C.D. and L.U. performed experiments and analyzed data. F.R., L.U. and J.H wrote the manuscript with the input from all authors.

