## Supplementary material for "Microglial P2X4 receptors promote ApoE degradation and cognitive deficits in Alzheimer disease": Sup data

Sup Fig 1

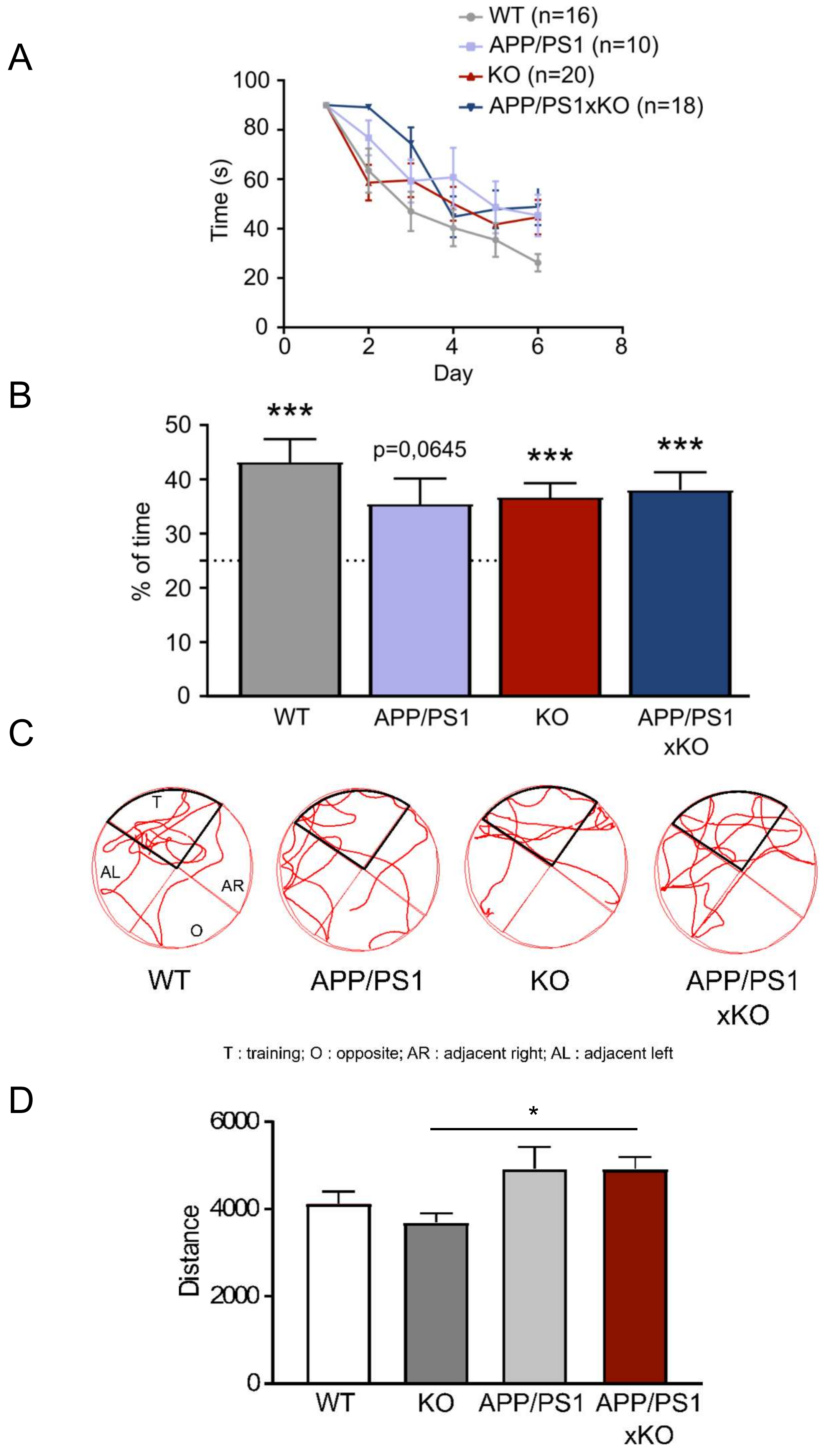
A
WT (n=16)
APP/PS1 (n=10)
KO (n=20)
APP/PS1xKO (n=18)
Time (s)
100
80
60
40
20
0
0
2
4
6
8
Day
B
***
p=0,0645
***
***
50
40
30
20
10
0
% of time
WT
APP/PS1
KO
APP/PS1
xKO
C
T
AL
AR
O
WT
APP/PS1
KO
APP/PS1
xKO
T : training; O : opposite; AR : adjacent right; AL : adjacent left
D
*
6000
4000
2000
0
Distance
WT
KO
APP/PS1
APP/PS1
xKO

Sup Fig 2

MKALWAVLLV TLLTGCLAEG EPEVTDQLEW QSNQPWEQAL NRFWDYLRWV
QTLSDQVQEE LQSSQVTQEL TALMEDTMTE VKAYKK**ELEE QLGPVAEETR**
ARLGKEVQAA QAR**LGADMED LR**NRLGQYRN EVHTMLGQST EEIRARLSTH
LRKMRKRLMR DAEDLQKRLA VYKAGAREGA ERGVSAIRER **LGPLVEQGR**Q
R**TANLGAGAA QPLR**DRAQAF GDRIR**GRLEE VGNQAR**DRLE EVREHMEEVR
SKMEEQTQQI R**LQAEIFQAR** LKGWFEPIVE DMHRQWANLM EK**IQASVATN**
**PIITPVAQEN Q**

Sup Fig 3

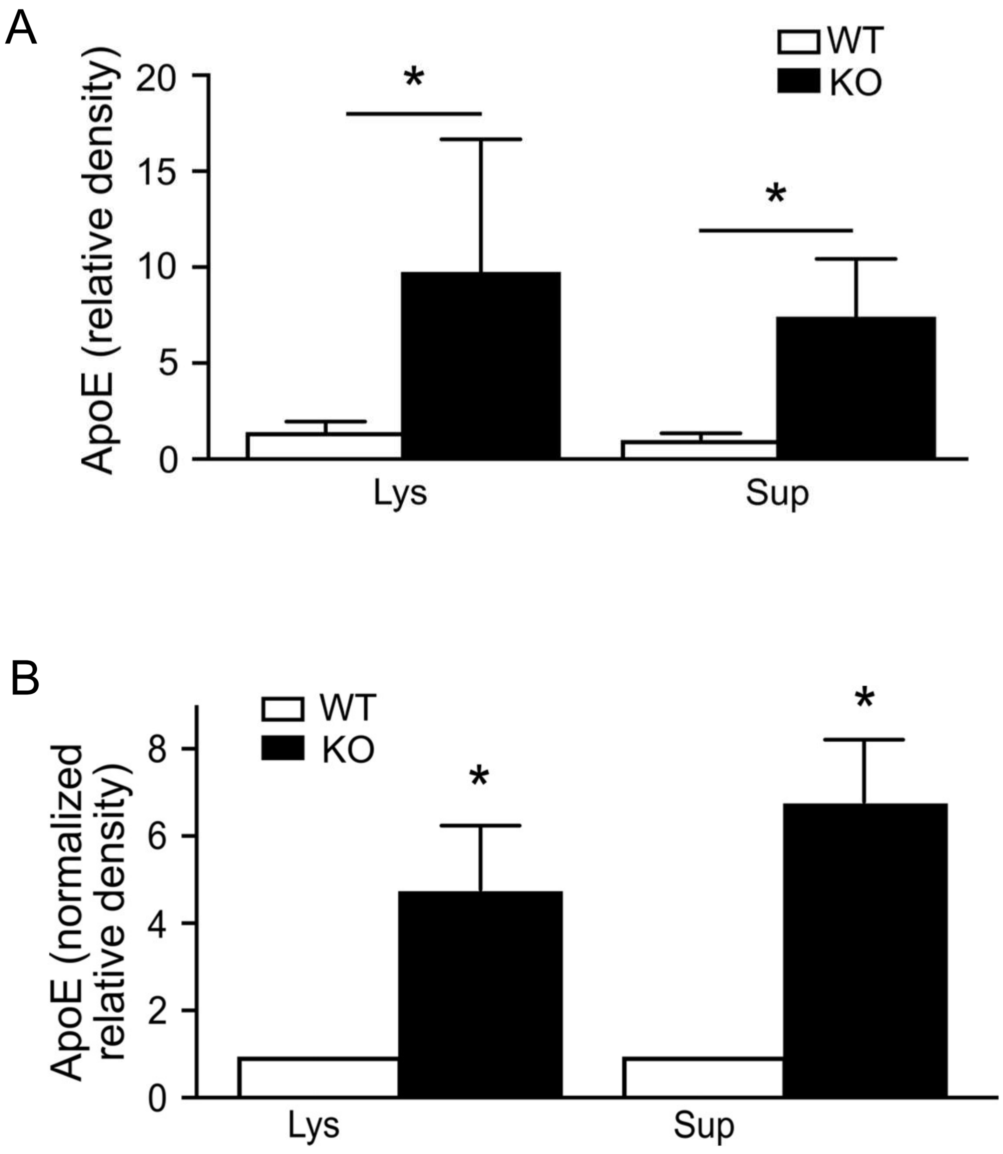
A
WT
KO
ApoE (relative density)
20
15
10
5
0
*
*
Lys
Sup
B
WT
KO
ApoE (normalized relative density)
8
6
4
2
0
*
*
Lys
Sup

Sup Fig 4

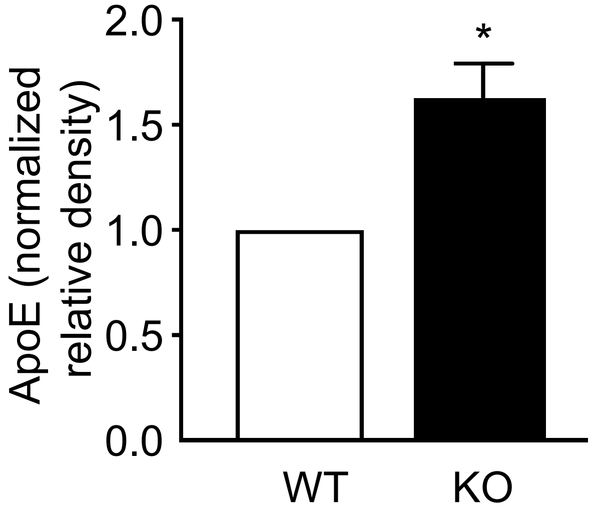

Sup Fig 5

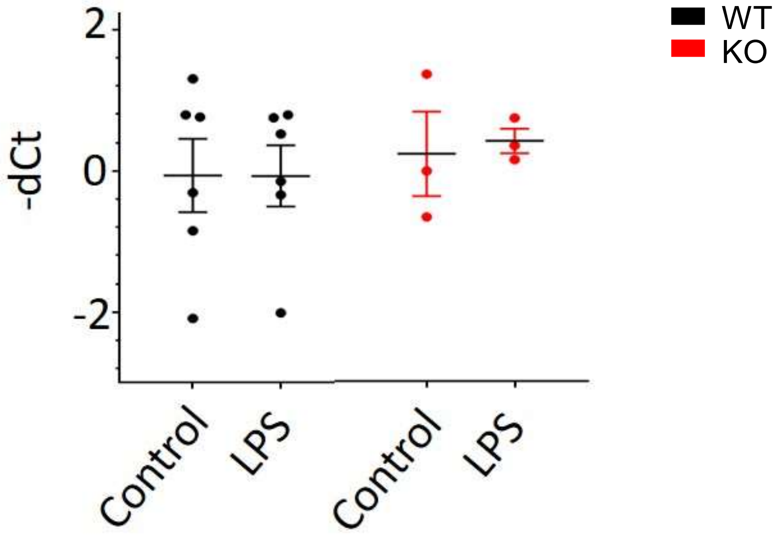

Sup Fig 6

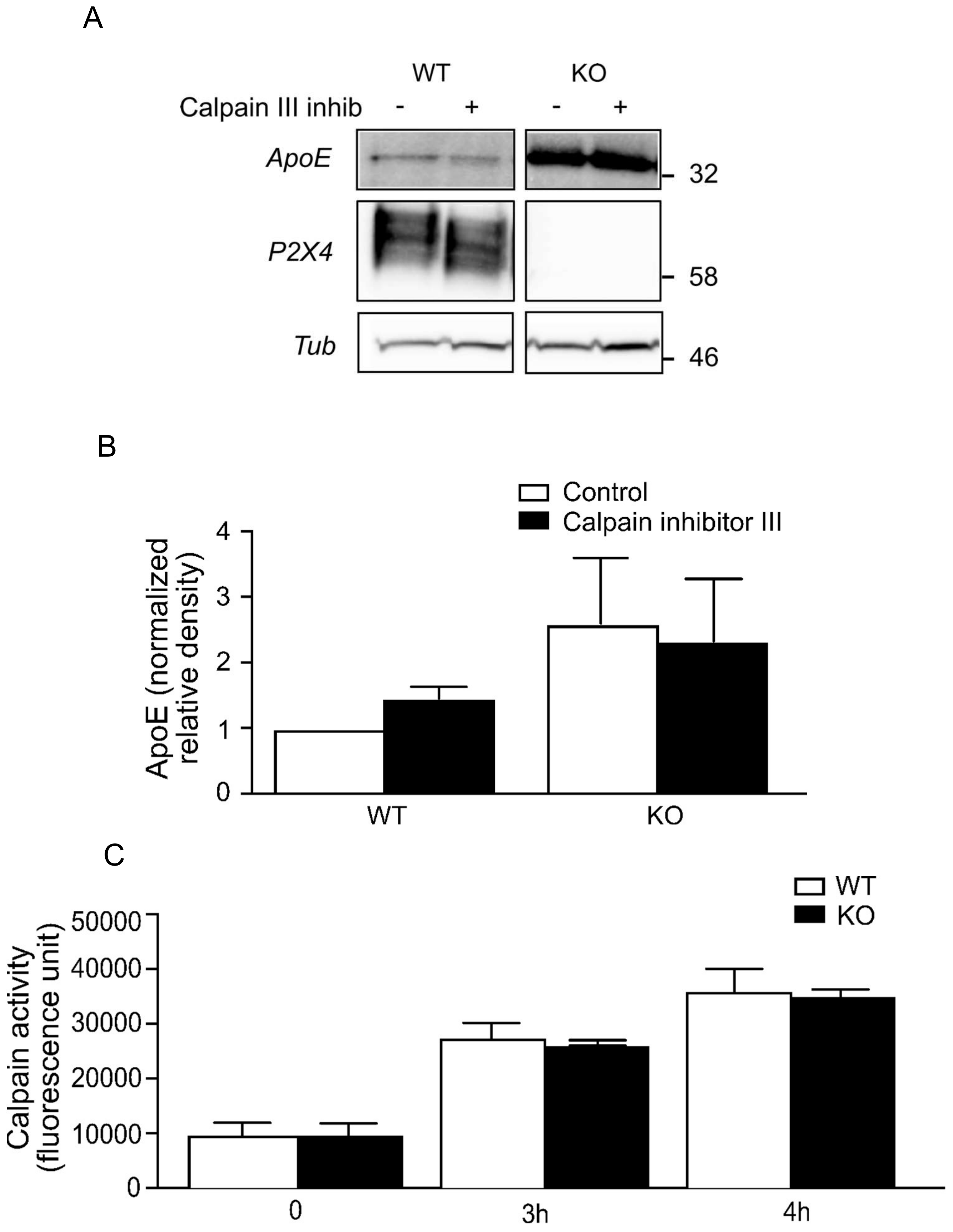
A
WT
KO
Calpain III inhib
-
+
-
+
ApoE
P2X4
Tub
32
58
46
B
Control
Calpain inhibitor III
ApoE (normalized relative density)
4
3
2
1
0
WT
KO
C
WT
KO
Calpain activity (fluorescence unit)
50000
40000
30000
20000
10000
0
0
3h
4h

Sup Fig 7

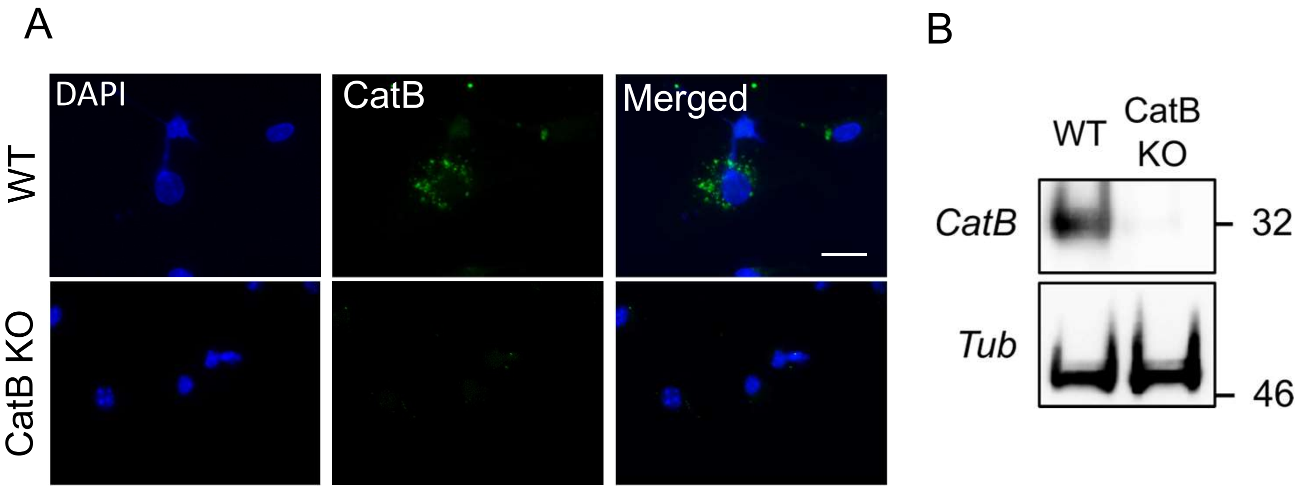
A
DAPI
CatB
Merged
WT
CatB KO
B
WT
CatB KO
CatB
32
Tub
46

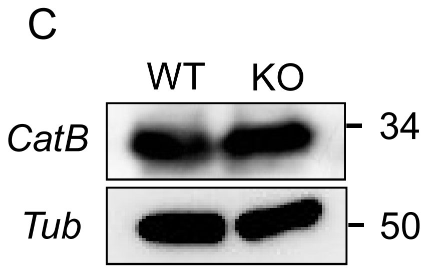
C
WT
KO
CatB
34
Tub
50

Sup Fig 8

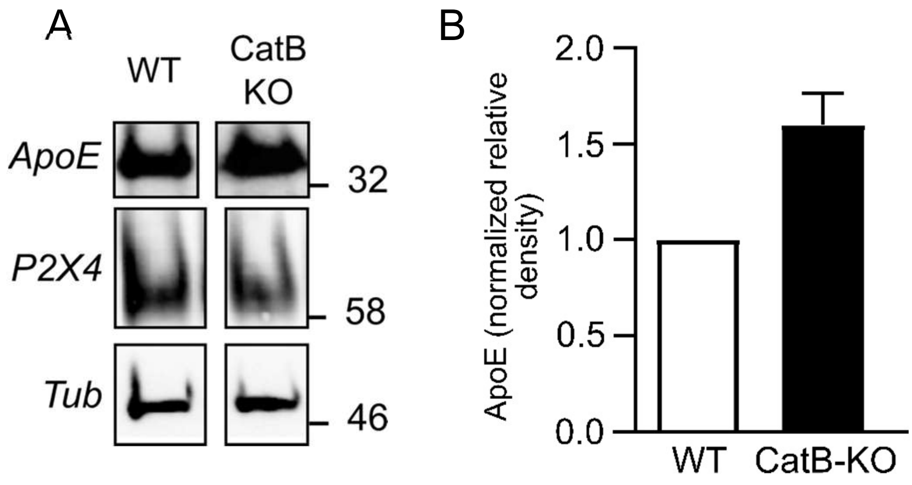

A
WT
CatB KO
ApoE
P2X4
Tub
32
58
46
B
2.0
1.5
1.0
0.5
0.0
ApoE (normalized relative density)
WT
CatB-KO

### Supplementary figure legends

**Sup figure 1: Increased memory performances of APP/PS1xKO mice in the Morris water maze.** (A) Comparison of the learning curve of WT, P2X4$^{-/-}$, APP/PS1 and APP/PS1xKO. The time spent to reach the platform for each day of training is reported for each genotype. Compared to WT, all other genotypes seem to learn slower but differences are not statistically different from WT, Two-way ANOVA. (B) Retention test analysis, 48h after the last day of training, time spent in the target quadrant was measured and compared to the theorical value of 25%. Only APP/PS1 mice show no statistical difference, whereas APP/PS1xKO spend significatively more time in the targeted quadrant. *** $p<0.001$, one sample Wilcoxon test. (C) Representative swim paths during the retention test for the 4 genotypes. (D) Activity of mice in the open field test. Total distance traveled by mice in the whole arena during 10 min recording. (n = 16, 10, 20 and 18 mice for WT, APP/PS1, KO and APP/PS1xKO respectively, * $p<0.05$, One-way ANOVA).

**Sup figure 2: Co-purification of Apolipoprotein E in the P2X4-Signaling Complex in mouse BMDM membrane extracts.** Membrane enriched fractions of mouse BMDM were immunoprecipitated with a specific anti-P2X4 antibody. Pulled-down protein complexes were separated by polyacrylamide gel electrophoresis, trypsinized and analyzed by LC MS/MS. Coverage of ApoE (Swiss-Prot P08226) by MS/MS identified peptides is indicated in green. Coverage results from two independent MS/MS experiments. Total coverage is 22.6% of ApoE sequence (84/371 residues).

**Sup figure 3: Comparison of non-normalized *versus* normalized ApoE levels in BMDM cells.** Comparison of ApoE western blot between non-normalized and normalized data. Amounts of ApoE in either lysates or supernatant of cultures is very variable, we therefore expressed the ratio of relative densities for KO over that of WT, and attributed an arbitrary value of 1 to WT, for statistical and graphical purposes . (A). Results from non-normalized experiments. Results are expressed as a ratio ApoE band density over that of tubulin obtained in lysates. (B). The same data as in A, normalized as explained above. Essentially similar results were obtained when data from the same experiments as in A were normalized to signal obtained in WT condition, albeit with lower variability. N = 4 independent cultures, * $p<0.05$, unpaired *t*-test.

**Sup figure 4**: Comparison of ApoE levels in WT and P2X4$^{-/-}$ microglial primary cultures. Western blot signal of ApoE was normalized to signal obtained from WT culture. As in BMDM, significant increase of ApoE is observed in P2X4$^{-/-}$ microglia cultures. n = 5 independent cultures, * p<0.05, unpaired *t*-test.

**Sup figure 5: Identical ApoE transcription in WT and P2X4$^{-/-}$ BMDM in control and LPS conditions.** BMDM from WT and P2X4$^{-/-}$ mice were stimulated or not with 100 ng of LPS overnight. After RNA extraction expression of ApoE mRNA was quantified by RT-qPCR. No difference in ApoE mRNA levels could be observed between the different conditions. n = 3 independent cultures, two-way ANOVA.

**Sup figure 6: Calpains are not involved in ApoE degradation**. (A) Representative western blot of ApoE from WT and P2X4$^{-/-}$ BMDM treated with 10 µM calpain inhibitor III. (B) Quantification of ApoE signal from A, normalized to WT condition n = 3 independent cultures, one sample Wilcoxon test compared to the theoretical value of 1. (C) Fluorescence of the fluorogenic calpain substrate Suc-leu-leu-val-tyr AMC (100 µM) was read after 3 h. No difference is observed between WT and KO BMDM cells.

**Sup figure 7: P2X4 deletion does not affect the expression of CatB.** (A, B) Validation of CatB antibody by immunocytochemistry and western blot. Scale bar 5 µm. (C) Western blot analysis of CatB expression in WT and KO BMDM cells shows no obvious difference of CatB expression between the two genotypes.

**Sup figure 8**: **Increase of ApoE in CatB KO BMDM.** (A) Representative western blot of ApoE from WT and CatB KO BMDM cell lysate. (B) Quantification of ApoE signal presented in A. n = 2 independent cultures.
